# N-cadherin facilitates trigeminal sensory neuron outgrowth and target tissue innervation

**DOI:** 10.1101/2024.05.20.594965

**Authors:** Caroline A. Halmi, Carrie E. Leonard, Alec T. McIntosh, Lisa A. Taneyhill

## Abstract

The trigeminal ganglion emerges from the condensation of two distinct precursor cell populations, cranial placodes and neural crest. While its dual cellular origin is well understood, the molecules underlying its formation remain relatively obscure. Trigeminal ganglion assembly is mediated, in part, by neural cadherin (N-cadherin), which is initially expressed by placodal neurons and required for their proper coalescence with neural crest cells. Axon outgrowth first occurs from placodal neurons, but as gangliogenesis proceeds, neural crest cells also differentiate into N-cadherin-expressing neurons, and both extend axons toward targets. However, the role of N-cadherin in axon outgrowth and target innervation has not been explored. Our data show that N-cadherin knockdown in chick trigeminal placode cells decreases trigeminal ganglion size, nerve growth, and target innervation *in vivo*, and reduces neurite complexity of neural crest-derived neurons *in vitro.* Furthermore, blocking N-cadherin-mediated adhesion prevents axon extension in most placodal neurons *in vitro*. Collectively, these findings reveal cell- and non-cell autonomous functions for N-cadherin, highlighting its critical role in mediating reciprocal interactions between neural crest- and placode-derived neurons throughout trigeminal ganglion development.

## INTRODUCTION

Sensory ganglia transmit various inputs from the body and environment to the central nervous system (Vermeiren et al., 2020). Within the head, the cranial sensory ganglia perform these functions for their respective cranial nerves. The trigeminal nerve, the largest and most complex of the cranial nerves, relays the majority of somatosensory information within the head (Heimer, 1983) through three major branches - the ophthalmic (OpV), maxillary (MxV), and mandibular (MnV) - that innervate the forehead and eye region, upper jaw, and lower jaw, respectively. The cell bodies for this nerve are positioned within the trigeminal ganglion, which arises from neural crest and placode cells (D’Amico-Martel and Noden, 1980; Hamburger, 1961). While both cell types differentiate into trigeminal sensory neurons (Hamburger, 1961), the neural crest also gives rise to glial cells. Trigeminal neural crest cells originate from the midbrain and anterior hindbrain (Lumsden et al., 1991; Noden, 1973), while the placodal contribution arises from the OpV and maxillomandibular (MmV) placodes. Although these placodes comprise the trigeminal placode to generate one structure, they differ in two important ways. First, they give rise to neurons to distinct lobes of the trigeminal ganglion (OpV versus MxV or MnV, respectively). Second, OpV placode cells begin exiting the cell cycle and differentiating into neurons as early as E1/ Hamburger-Hamilton stage (HH)8 and continue until E3.5/HH21 (McCabe et al., 2009), traveling to the trigeminal ganglion-forming region as a combination of neuroblasts and differentiated neurons. Conversely, the MmV placode generates neuroblasts that do not differentiate or become post-mitotic until they reach the ganglionic anlage (Begbie, 2002; Graham and Shimeld, 2013; McCabe et al., 2009; Shiau et al., 2011). Notably, in the chick, placodal neurons are the first neurons present within the trigeminal ganglion, differentiating and growing towards the periphery to make initial connections with their targets within 2-5 days of development. In contrast, neural crest cells only begin neuronal differentiation at 4 days of development, and continue differentiating up to 7 days (D’Amico-Martel and Noden, 1983, 1980).

Reciprocal interactions between neural crest cells and placodal neurons are required to properly form the trigeminal ganglion (Hamburger, 1961; Moody and Heaton, 1983; Shiau et al., 2008). Much of this research has focused on the influence of neural crest cells on surrounding cell types, but less is known about how placode cells impact the neural crest. Because placodal neurons are present prior to neural crest-derived neurons, they may serve as pioneer neurons, providing a scaffold for later-born neurons to grow along as they innervate their targets (Bate, 1976). Although supported by some studies (Moody et al., 1989; Stainier and Gilbert, 1990), additional evidence for this is needed, and the molecules orchestrating this process remain unknown.

Intercellular adhesion among placode cells and neural crest cells regulates trigeminal gangliogenesis (Steventon et al., 2014) and may also facilitate trigeminal axon outgrowth to targets, with cadherins promoting adhesion between adjacent cells and acting as mediators of gangliogenesis (Halmi et al., 2022; Shiau and Bronner-Fraser, 2009; Theveneau et al., 2013; Wu and Taneyhill, 2019). The cadherin extracellular domain facilitates *cis* and *trans* interactions via homophilic and heterophilic binding (Basu et al., 2017; Halmi et al., 2022; Straub et al., 2011) while the intracellular domain contains two binding sites for catenin proteins (p120-catenin and β-catenin) that regulate cadherin adhesive ability and provide a link to the actin cytoskeleton, respectively (Niessen et al., 2011). Neural cadherin (N-cadherin) is expressed in neural tissue at various stages of development (Hatta and Takeichi, 1986). It is also found in mesodermal (Dady et al., 2012; Nakagawa and Takeichi, 1997) and ectodermal derivatives, including trigeminal and epibranchial placode cells prior to their delamination (Shiau and Bronner-Fraser, 2009), and later in all trigeminal sensory neurons (Inuzuka et al., 1991; Itoh et al., 1997; Redies et al., 1992; Tan et al., 2010). While multiple studies, largely in the central nervous system, describe roles for N-cadherin in regulating neurite formation (Bixby and Zhang, 1990; Gärtner et al., 2012; Hansen et al., 2008), axon fasciculation (Masai et al., 2003), and synapse formation (Benson and Tanaka, 1998; Jontes, 2018; Riehl et al., 1996), the function of N-cadherin in sensory neurons has not been thoroughly characterized. Although N-cadherin function in placode cells is critical for the early condensation of the chick trigeminal ganglion from undifferentiated neural crest cells and placodal neurons (Shiau and Bronner-Fraser, 2009), whether N-cadherin mediates the later growth and maturation of all trigeminal ganglion sensory neurons, and specifically neurite production and axon extension, is unknown. Furthermore, how N-cadherin-expressing placodal neurons impact the development of later-born neural crest-derived neurons is poorly understood.

Building upon these prior studies, we demonstrate that placodal expression of N-cadherin is vital for later chick trigeminal neurodevelopment. We observed substantial deficits in trigeminal ganglion morphology and axonal projections to target tissues that arise from effects on both placode- and neural crest-derived neurons in response to N-cadherin depletion from placodal neurons. These latter defects are noted both *in vivo* and *in vitro,* providing evidence to support the hypothesis that N-cadherin-mediated adhesion is important for trigeminal neurodevelopment. Strikingly, we note decreased neurite production in neural crest-derived neurons that correlates with altered β-catenin levels, pointing to non-cell autonomous effects on the neural crest after N-cadherin knockdown in placodal neurons. These results indicate that trigeminal neurons depend on the adhesive function of N-cadherin to facilitate the outgrowth of axons. Altogether, our findings reveal a new long-term role for N-cadherin in driving the proper outgrowth of both placode- and neural crest-derived neurons throughout trigeminal gangliogenesis.

## RESULTS

### N-cadherin knockdown in placode cells disrupts trigeminal ganglion development

To elucidate the function of N-cadherin in placodal neurons during the formation and maturation of the trigeminal ganglion, a validated N-cadherin morpholino (Shiau and Bronner-Fraser, 2009) (MO) was unilaterally electroporated into chick trigeminal placode cells at a single timepoint prior to their delamination. Embryos were grown to various endpoints (2.5-6.5 days post-incubation) to assess trigeminal ganglion morphology. Initial experiments with a 5 base pair mismatch N-cadherin control MO (N-cadherin MM) (Shiau and Bronner-Fraser, 2009) confirmed that the electroporation technique did not cause any morphological changes (Fig. S1). Once validated, the unelectroporated (contralateral) sides of N-cadherin MO-electroporated embryos were used as controls for all *in vivo* experiments to account for any developmental variation. Trigeminal ganglion size and nerve morphology were analyzed by whole-mount immunohistochemistry using an antibody against tubulin beta 3, class III (Tubb3) to label neurons, followed by tissue clearing for imaging. Given the time difference in neuronal differentiation in the trigeminal ganglion (D’Amico-Martel and Noden, 1983; Moody et al., 1989), the Tubb3 antibody was presumed to identify placode-derived neurons up to E4/HH23, after which both placode- and neural crest-derived neurons were labeled indistinguishably.

During initial gangliogenesis (E2.5/HH16-18), N-cadherin depletion from trigeminal placode cells caused neuron dispersal, increasing the forming trigeminal ganglion area compared to the contralateral control (Fig. 1A-A’, 1F), consistent with previous work (Shiau and Bronner-Fraser, 2009). However, by E3.5/HH19-22, the size of the trigeminal ganglion on the electroporated side was significantly reduced compared to the contralateral control (Fig. 1B-B’, 1F). This reduction persisted even as neural crest cells began differentiating into neurons at E4.5/HH23-25 (Fig. 1C-C’, 1F), pointing to potential effects on both placode- and neural crest-derived neurons during trigeminal neurodevelopment. At both E5.5/HH26-27 and E6.5/HH28-30, N-cadherin MO-electroporated trigeminal ganglia were no longer statistically different from controls (Fig. 1D-E’, 1F).

**Figure 1.**
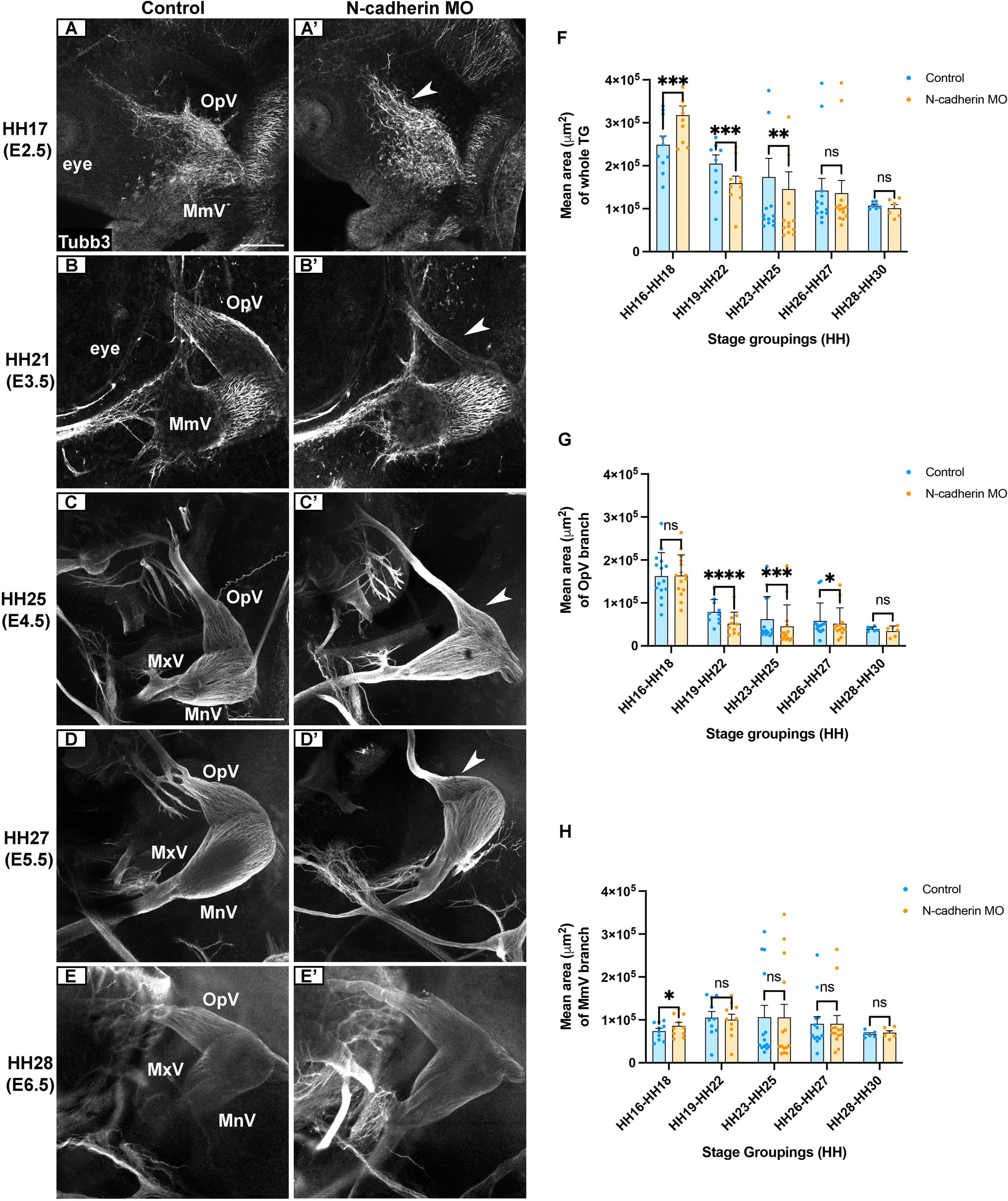
Knockdown of N-cadherin in placode cells disrupts trigeminal ganglion development. (A-E’) Representative images of the trigeminal ganglion (contralateral control and N-cadherin MO-electroporated side) after Tubb3 immunostaining (E2.5/HH16-18: A-A’ (n=10), E3.5/HH19-22: B-B’ (n=10), E4.5/HH23-25: C-C’ (n=14), E5.5/HH26-27: D-D’ (n=13), E6.5/HH28-30: E-E’ (n=6)). Arrowheads highlight reductions in OpV lobe size in N-cadherin MO samples compared to controls. Scale bar in (A) is 250 µm and applies to (A’-B’); scale bar in (C) is 250 µm and applies to (C’-E’). (F-H) Mean area ± SEM of whole trigeminal ganglia (F), OpV branch (G), and MmV/MxV branch (H) for contralateral and N-cadherin MO sides of embryos. Statistical significance was determined via paired t-tests and Wilcoxon signed-rank test. (*) p<0.05, (**) p<0.01, (***) p<0.001, (****) p<0.0001. Abbreviations: TG: trigeminal ganglion; OpV: ophthalmic; MmV: maxillomandibular; MxV: maxillary; MnV: mandibular; ns: not significant.

Next, we measured the area of OpV and MmV individually. Interestingly, N-cadherin knockdown led to a significant increase in MmV, but not OpV, at E2.5/HH16-18 (Fig. 1G-H). However, from E3.5-E6.5/HH19-30, N-cadherin depletion did not affect MmV lobe development (Fig. 1H) but led to a significant reduction in OpV lobe size (Fig. 1D-D’, 1G). By E6.5/HH28-30, the size of both lobes was similar to controls (Fig. 1G-H). We did not observe any increased cell proliferation in N-cadherin MO OpV lobes at E6.5/HH28-30, as indicated by phospho-histone H3 expression (Fig. S2), that would explain the absence of phenotype observed at this stage.

To investigate whether specific subpopulations of trigeminal sensory neurons were being impacted after N-cadherin knockdown, we analyzed TrkA and TrkB receptor expression in OpV sections at E5.5/HH26-27 when both placode- and neural crest-derived neurons are present in the trigeminal ganglion, as this stage grouping include neural crest cell differentiation into neurons (D’Amico-Martel and Noden, 1983; Williams et al., 1995). TrkA and TrkB neurons were counted on immunolabeled sections from contralateral (Fig. S3A-B’) and N-cadherin MO (Fig. S3C-D’) trigeminal ganglia. N-cadherin MO treatment caused a significant decrease in TrkB-positive neurons (Fig. S3E); however, after normalization against the smaller area of the OpV lobe, the TrkB neuron count was no longer statistically different from controls (Fig. S3F). No changes were observed in TrkA-positive neurons (Fig. S3E-F). These results suggest that the morphological deficits observed in the trigeminal ganglion over time cannot be attributed to alterations in specific neuronal subpopulations.

### Depletion of N-cadherin from trigeminal placode cells disrupts target innervation by OpV nerves

Since extensive morphology changes were observed within OpV (Fig. 1G-H), we focused subsequent *in vivo* analyses on its development and next evaluated whether OpV axons and nerve projections into cranial target tissues were also affected. We first examined the lateral nasal branch at E4.5/HH23-25, as it becomes a prominent structure within the embryonic head by this stage (Higashiyama and Kuratani, 2014). We performed whole-mount Tubb3 immunohistochemistry, which labels both placode- and neural crest-derived neurons at this stage, and compared the N-cadherin MO-electroporated side of the embryo to the contralateral control side. These experiments revealed fewer lateral nasal nerve branches on the N-cadherin MO side of the embryo (Fig. 2A-B). To quantify these findings, we assessed the distribution of terminal nerve endings at 30 µm intervals, starting at the first axon branching point from the OpV lobe. No difference in the number of terminal nerve endings was observed closer to the trigeminal ganglion, but fewer were present in distal regions after N-cadherin MO treatment (Fig. 2C). Moreover, nerve width and innervation area were significantly reduced after N-cadherin knockdown compared to contralateral control trigeminal ganglia (Fig. 2D-E).

**Figure 2.**
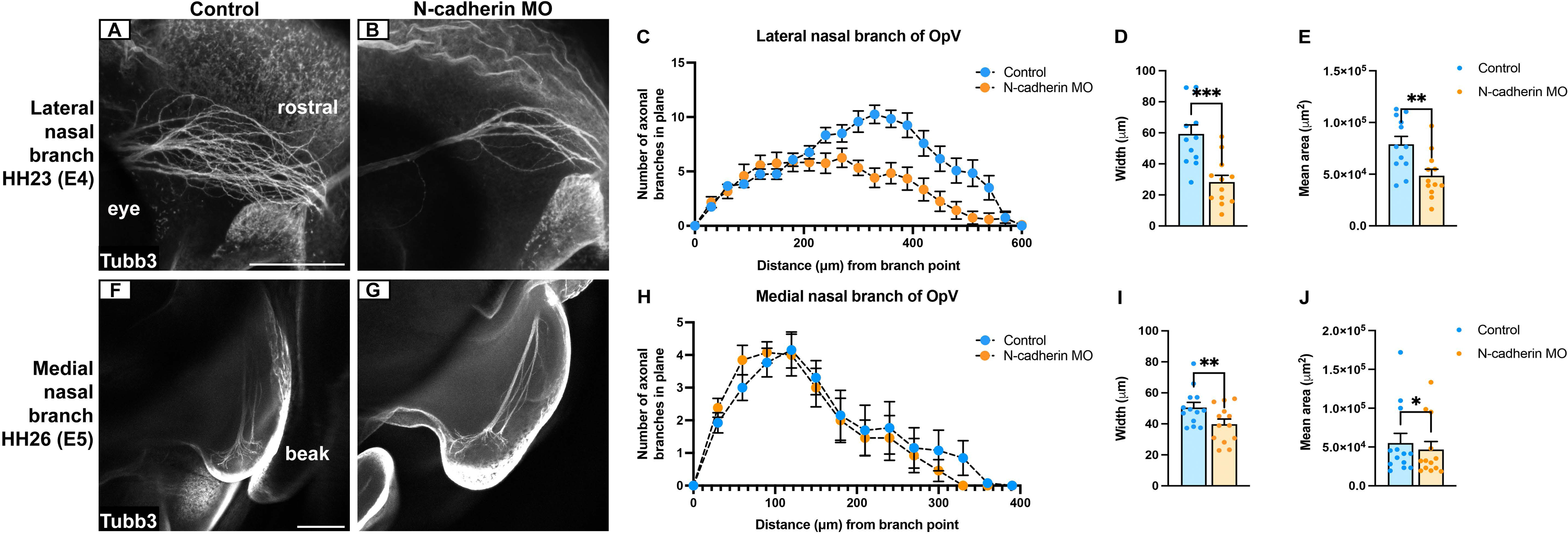
N-cadherin depletion from placode cells negatively impacts outgrowth and target tissue innervation for OpV nerve branches. (A, B, F, G) Representative images of the lateral (A, B) and medial (F, G) nasal nerves (contralateral control and N-cadherin MO-electroporated side) after Tubb3 immunostaining (A, B: E4.5/HH23-25 (n=12); F, G: E5.5/HH26-27 (n=13)). Scale bars in (A) and (F) are 250 µm and apply to (B) and (G), respectively. (C, H) Mean ± SEM of terminal nerve endings. Significant distances in (C) are observed at 420, 450, 480, and 530 µm (*); 300 and 570 µm (**); and 330, 360, and 390 µm (***). No significant differences were observed in (H). (D, I) Mean nerve width ± SEM. (E, J) Mean innervation area ± SEM. Statistical significance was determined via paired t-tests (C-E, H-J), with adjustments made for multiple comparisons (C, H). (*) p<0.05, (**) p<0.01, (***) p<0.001.

Using the same method, we evaluated the medial nasal nerve at E5.5/HH26-27, as this stage provided more consistent morphological development compared to E4.5/HH23-25 embryos (Higashiyama and Kuratani, 2014), where increased variability was observed. While the number of medial nasal terminal nerve endings was similar between conditions (Fig. 2F-H), the nerve width and target innervation area were significantly smaller after N-cadherin knockdown compared to contralateral control trigeminal ganglia (Fig. 2I-J). Together, these findings reveal that N-cadherin is required by OpV trigeminal neurons to send axons to targets, thus leading to proper nerve outgrowth, fasciculation, and target innervation.

### N-cadherin knockdown in placode cells leads to increased cell death within the OpV lobe

To assess whether cell death altered OpV lobe size, axon outgrowth, and target innervation after N-cadherin knockdown in placode cells, we performed staining for apoptotic cells using cleaved caspase-3 and TUNEL (Fig. 3). We examined timepoints encompassing when placode cells are first delaminating (E1.5/HH13-15) and when morphological changes were observed (Fig. 1: E2.5/HH16-18, E3.5/HH19-22, E4.5/HH23-25). Apoptosis was evident in tissue sections from unelectroporated, contralateral control (Fig. 3A-A’) and N-cadherin MO-electroporated (Fig. 3B-B’) trigeminal ganglia. When correcting for size differences in the ganglion after N-cadherin knockdown, cell death was significantly increased on the N-cadherin MO-treated side only during initial coalescence of placodal neurons with undifferentiated neural crest cells (E2.5/HH16-18), when OpV size remained unchanged (Fig. 1A-A’, 1G). Once placodal neurons and neural crest cells have formed the trigeminal ganglion, and neural crest cells have begun differentiating into neurons beyond E4, there was no significant difference in cell death between the N-cadherin MO-treated and contralateral control sides (Fig. 3C), despite reduced OpV size at these stages after N-cadherin knockdown (Fig. 1B-D’, 1G). These data suggest that apoptosis could partially account for some of the morphological phenotypes observed during early gangliogenesis prior to neural crest cell differentiation into trigeminal sensory neurons.

**Figure 3.**
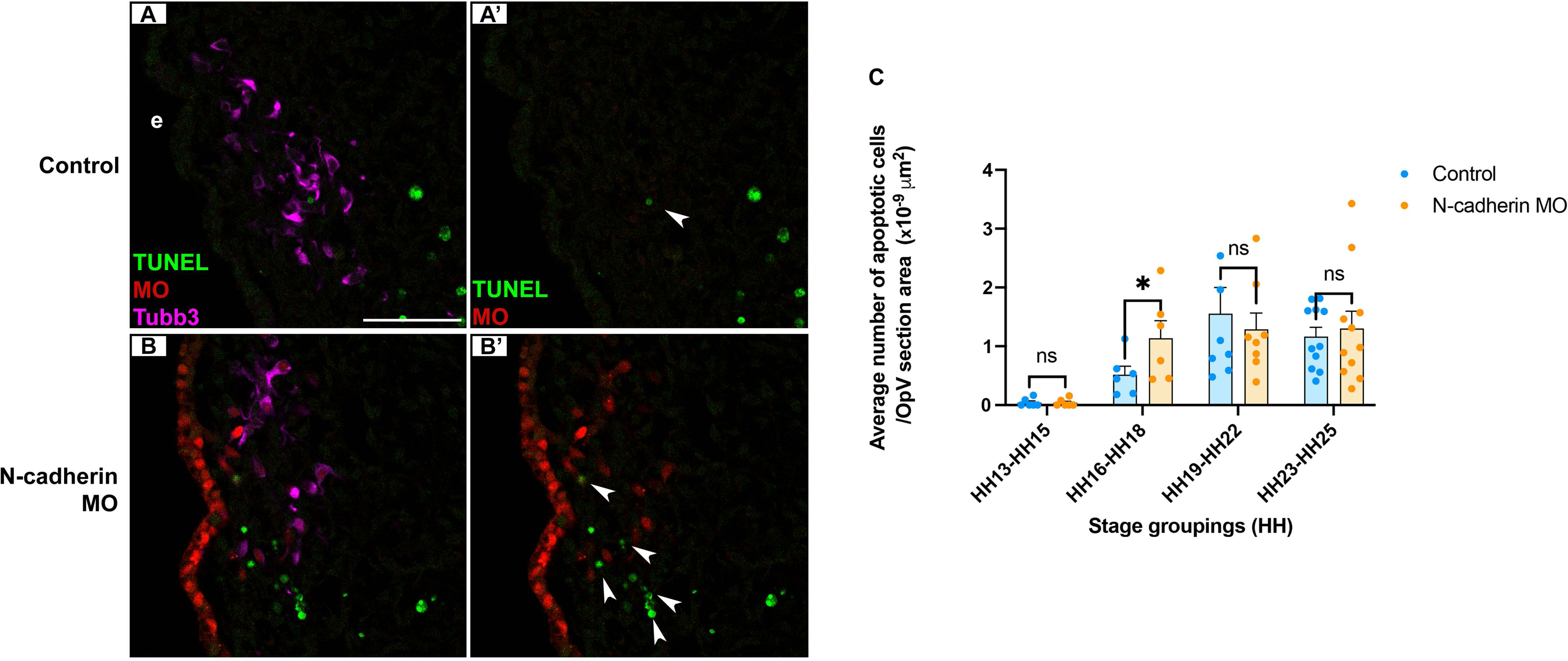
Morphological changes in the trigeminal ganglion after N-cadherin knockdown in placode cells are in part due to increased cell death. (A-B’) Transverse sections through the OpV lobe (contralateral control and N-cadherin MO-electroporated side) from representative E2.25/HH16 embryos with Tubb3 immunostaining and TUNEL staining. Scale bar in (A) is 50 µm and applies to all images. (C) Five serial sections per embryo from stage groupings E1.5/HH13-16 (n=6), E2.5/HH16-18 (n=6), E3.5/HH19-22 (n=8), and E4.5/HH23-25 (n=11) were imaged and used for analysis. Arrowheads in (A’, B’) mark TUNEL puncta. Statistical significance was determined via paired t-tests and Wilcoxon signed-rank test. (*) p<0.05. Abbreviations: e: ectoderm; ns: not significant.

### Neurite complexity in neural crest-derived neurons is reduced after N-cadherin knockdown in placode cells

The absence of persistent apoptosis and presence of continued axon outgrowth deficits, particularly at stages when neural crest cells differentiate into neurons, led us to hypothesize that depletion of N-cadherin from placode cells may be impacting later-born neural crest-derived neurons. To test this, we performed a sequential electroporation (Halmi et al., 2022) using a genome-incorporating vector (PiggyBacGFP) and its accompanying transposase (PBase) (Lu et al., 2009; Macdonald et al., 2012) to first label neural crest cells with a stably expressed green fluorescent protein (GFP), followed by electroporation of placode cells with N-cadherin MO or the validated N-cadherin mismatch (MM) control morpholino (Fig. S1), which was used as the control for this experiment (Shiau and Bronner-Fraser, 2009). Sequential electroporations of neural crest cells and placode cells leads to mutually exclusive expression of the introduced plasmid/morpholino (Fig. S4), as shown previously (Halmi et al., 2022). Sequentially-labeled trigeminal ganglia were dissected, cultured as explants, grown *in vitro* for 3 days, and subsequently processed for immunostaining with GFP and Tubb3 antibodies (Fig. 4A). Although N-cadherin MM-treated explant cultures exhibited numerous axons extending from neural crest-derived neurons (Fig. S5A-A’), qualitatively we observed relatively few GFP-positive neural crest-derived axons in N-cadherin MO-treated samples (Fig. S5B-B’). Interestingly, neurites, which precede axon specification and extension (Sainath and Gallo, 2015), were noted from GFP-positive neural crest-derived somata in both N-cadherin MO- and N-cadherin MM-treated explant cultures. These were numerous in N-cadherin MM-treated explant cultures (Fig. 4B-C), but noticeably fewer in N-cadherin MO-treated explants (Fig. 4D-E). In addition to a significant decrease in the number of primary and secondary neurites in neural crest-derived neurons, tertiary neurites were absent in N-cadherin MO-treated samples but present in N-cadherin MM-treated explants (Fig. 4F). Although neurite branching was deficient in N-cadherin MO-treated explants cultures, there was no difference in the length of primary and secondary neurites compared to N-cadherin MM-treated explant cultures (Fig. 4G). These results indicate that N-cadherin in placodal neurons non-cell autonomously facilitate axon extension and branching in neural crest-derived neurons *in vitro*, providing an additional explanation for the sustained morphological deficits noted in the trigeminal ganglion *in vivo*.

**Figure 4.**
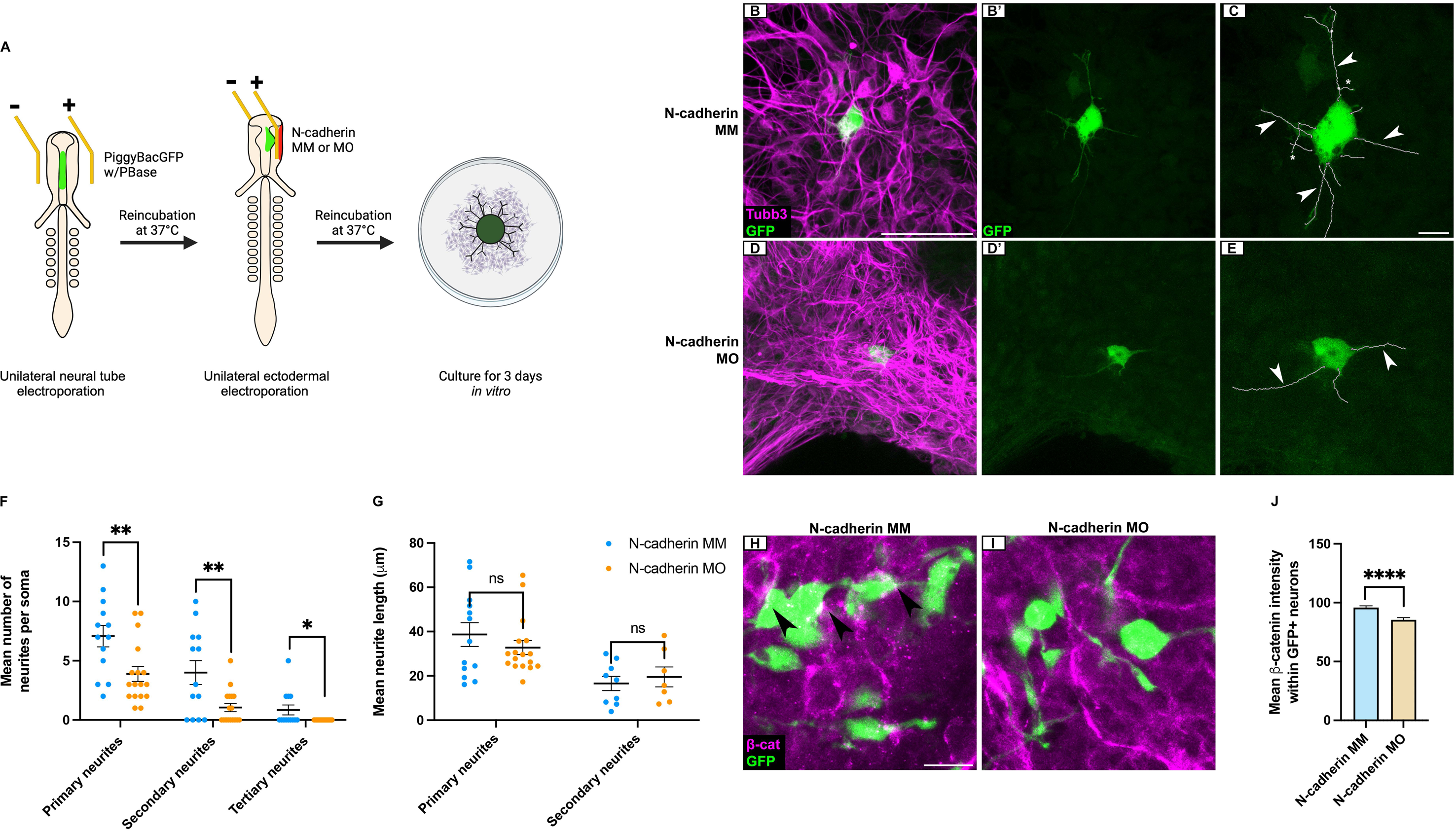
N-cadherin depletion from placode cells decreases neurite complexity in neural-crest derived neurons. (A) Schematic of sequential electroporation explant culture technique. (B-E) Representative maximum intensity Z-projections of GFP-labeled neural crest-derived neurons after placodal electroporation with N-cadherin MM (B-B’, control, (n=13)) or N-cadherin MO (D-D’, (n=17)) from trigeminal ganglion explants following GFP and Tubb3 immunostaining, and neurite-traced images (C, E). Arrowheads and asterisks denote examples of primary and secondary neurites, respectively. Scale bar in (B) is 50 µm and applies to (B’, D-D’); scale bar in (C) is 10 µm and applies to (E). Mean number (F) and length (G) ± SEM of GFP-positive neural crest-derived neurites after N-cadherin MM or N-cadherin MO electroporation. (H-I) Representative maximum Z-projections of GFP-labeled neural crest-derived neurons after placodal electroporation with N-cadherin MM (H, n=3) or N-cadherin MO (I, n=4) from trigeminal ganglion explants following GFP and β-catenin immunostaining. Arrowheads indicate colocalization of GFP and β-catenin staining. Scale bar in (H) is 10 µm and applies to (I). (J) Mean β-catenin signal intensity ± SEM in GFP-positive neural crest-derived neurons. Statistical significance was determined via unpaired t-tests and Mann-Whitney tests (F-G). (*) p<0.05, (**) p<0.01, (****) p<0.0001. Abbreviations: ns: not significant.

To explore a potential mechanism behind this observation, we examined β-catenin within neural crest-derived neurons in this same assay, given its known interactions with N-cadherin and role in neurite outgrowth (Benson and Tanaka, 1998; Halbleib and Nelson, 2006; Miyamoto et al., 2015). While membranous β-catenin was evident in neural crest-derived neurons of N-cadherin MM-treated samples (Fig. 4H, arrowheads), β-catenin levels were significantly reduced in neural crest-derived neurons of N-cadherin MO-treated samples, with less β-catenin evident along cell membranes (Fig. 4I, 4J). These findings support a mechanism whereby changes to β-catenin levels could contribute to the defects observed in neural crest-derived neurite production after N-cadherin knockdown in placode cells.

### N-cadherin controls proper axon outgrowth of trigeminal neurons *in vitro*

The morphological defects observed after N-cadherin knockdown in placode cells *in vivo* and *in vitro* suggest a role for N-cadherin-mediated adhesion in facilitating axon outgrowth of both placode- and neural crest-derived trigeminal sensory neurons. To test this, we utilized a validated function-blocking antibody against the extracellular domain of N-cadherin, known as MNCD2 (Bronner-Fraser et al., 1992; Redies et al., 1992) (anti-N-cadherin), to block N-cadherin-mediated adhesion in explanted trigeminal ganglion tissue.

Trigeminal ganglion explant cultures were grown *in vitro* for 24 hours, followed by treatment with anti-N-cadherin or a control rat IgG antibody for an additional 24 hours (Fig. 5A), and then processed for Tubb3 immunostaining (Fig. 5B, C, D). Wildtype explants incubated *in vitro* for 24 hours without treatment served as controls (“pre-treatment”) to evaluate baseline axon outgrowth and the possibility of axon pruning prior to antibody treatment. Ridge detection was used to detect segments, which were used as a proxy for individual axons (Fig. 5B’, C’, D’), and an intersection analysis was run to measure changes to axon outgrowth and branching (Fig. 5B”, 5C”, D”). While the Tubb3 fluorescence intensity between pre-treatment explants and anti-N-cadherin-treated samples was similar, a 60% reduction in Tubb3 fluorescence was observed in anti-N-cadherin-treated samples compared to rat IgG-treated samples (Fig. 5E). These results point to decreased axon density in trigeminal ganglion explant cultures after blocking N-cadherin-based adhesion. Furthermore, intersection analyses revealed that explants treated with anti-N-cadherin exhibited a significant increase in axonal branching compared to pre-treatment samples up to 1400 µm, while rat IgG-treated explants exhibited a significant increase in branching up to 1750 µm compared to the pre-treatment group (Fig. 5F). However, when comparing the anti-N-cadherin- and rat IgG-treated explants, significant differences were only observed up to 700 µm (Fig. 5F). Interestingly, of the neurons that did extend axons, the mean segment length of detected axons was similar between pre-treatment and anti-N-cadherin-treated explants (Fig. 5G), suggesting that some axons can extend in the absence of N-cadherin-mediated adhesion. However, the rat IgG-treated group had a significantly decreased average segment length compared to both, providing further support for increased axon branching over time. To verify this and confirm that the decreased segment length was not due to axon pruning, we visualized all detected segments using a violin plot to assess the distribution of the data (Fig. 5H). This analysis confirmed an increase in smaller segments in the rat IgG-treated group, validating that the observed decrease in mean segment length (Fig. 5G) resulted from averaging long and short segments, rather than axons retracting over time. Collectively, these data suggest N-cadherin-mediated adhesion promotes axonal outgrowth and complexity in trigeminal neurons *in vitro* but does not likely function in axon trimming.

**Figure 5.**
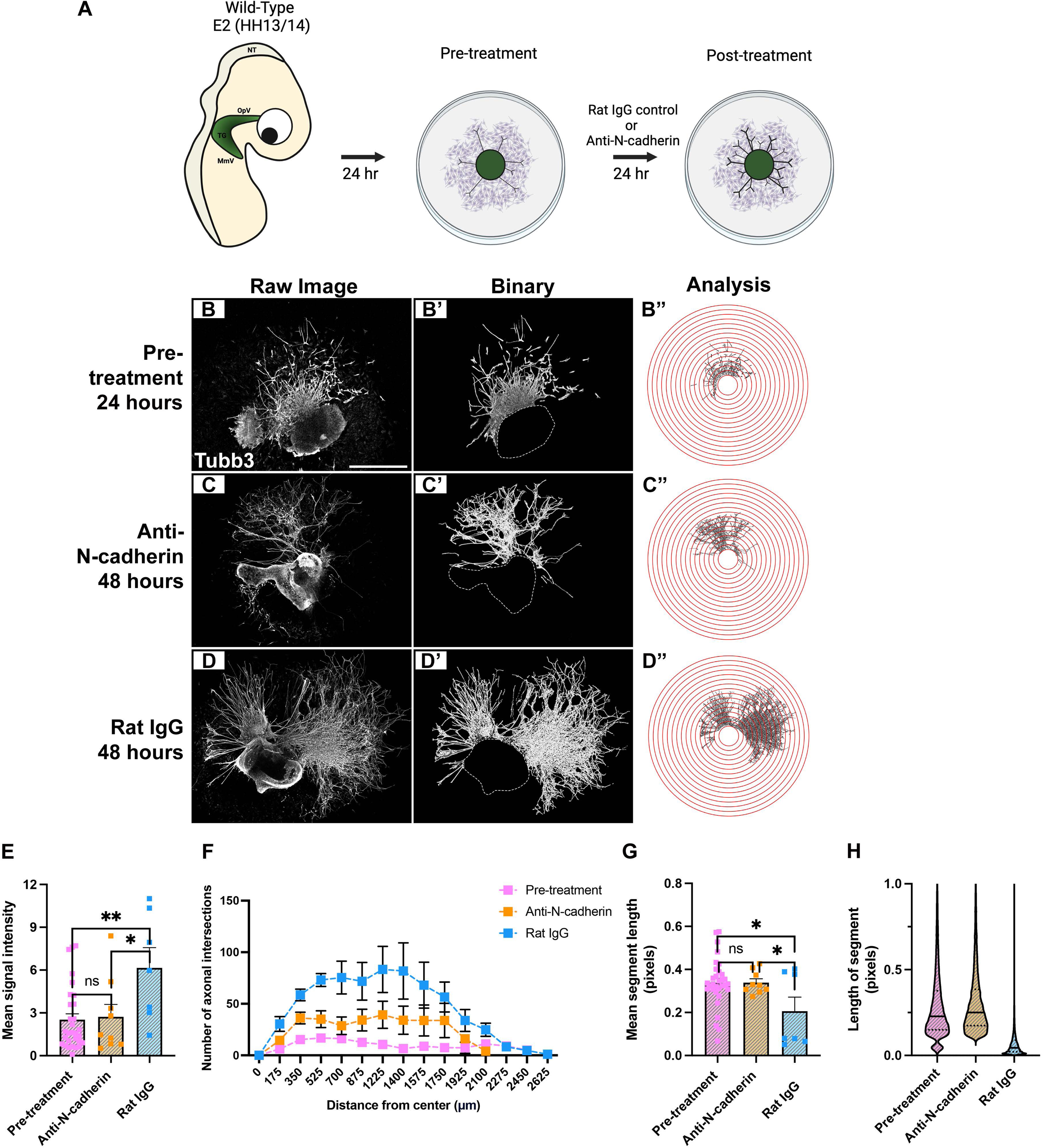
Blocking N-cadherin-mediated adhesion reduces trigeminal neuron axon outgrowth. (A) Schematic of N-cadherin function-blocking assay. (B-D”) Workflow for *in vitro* analyses. (B, C, D) Representative images of wildtype trigeminal ganglion explants after 24 hours (hr) of incubation and fixation (pre-treatment control explants (n=31)), or post-treatment with either rat IgG (n=7) or anti-N-cadherin (n=9) antibody, followed by Tubb3 immunostaining. Scale bar in (B) is 1 mm and applies to (C, D). (B’, C’, D’) Binary images generated after ridge detection processing on Fiji for pre-treatment, anti-N-cadherin-treated, or rat IgG-treated explants, respectively. Dotted lines indicate perimeter of explanted tissue cropped during processing. (B”, C”, D”) Processed images overlaid with concentric circle function to count the number of axonal intersections at increasing intervals of 175 µm. (E) Mean signal intensity for explant outgrowth ± SEM. (F) Mean number of axonal intersections ± SEM. Significant differences between pre-treatment control explants and anti-N-cadherin-treated explants are observed at 525 µm (*) and 175, 350, 875, 1225, and 1400 µm (**). Significant differences between pre-treatment controls and rat IgG-treated explants are observed at 1750 µm (*), 1575 µm (**), and 175-1400 µm (****). Significant differences between anti-N-cadherin- and rat IgG-treated explants are observed at 175, 350, and 700 µm (*) and 525 µm (**). (G) Mean segment length ± SEM. (H) Violin plots of segment lengths for pre-treatment, anti-N-cadherin-treated, and rat IgG-treated explants. Solid black lines indicate median and dotted black lines indicate interquartile range. Statistical significance was determined via unpaired t-tests and Mann-Whitney tests (E-G) with adjustments made for multiple comparisons (F). (*) p<0.05, (**) p<0.01, (***) p<0.001, (****) p<0.0001.

### N-cadherin is required cell autonomously for the axon outgrowth of the majority of placodal neurons

Our preceding findings indicate that most trigeminal sensory neurons rely on N-cadherin for axon outgrowth. To investigate their cellular origin, an ectodermal co-electroporation with a stably expressed red fluorescent protein (PiggyBacRFP plus PBase) (Lu et al., 2009; Macdonald et al., 2012) was performed to label placode-derived neurons. Next, RFP-positive explant cultures were treated with either rat IgG or anti-N-cadherin as previously described. Following immunostaining with Tubb3 and RFP antibodies, robust axon outgrowth was observed, the majority of which overlapped with RFP-positive placode-derived neurons after rat IgG treatment (Fig. 6A). Many Tubb3-positive projections were also RFP-positive in these control explant cultures (Fig. 6B-B”). Explant cultures treated with anti-N-cadherin, however, exhibited reduced overall axon outgrowth and fewer RFP-positive projections (Fig. 6C), as noted previously (Fig. 5C). While colocalization of Tubb3 and RFP was maintained in the anti-N-cadherin-treated axonal projections, the total amount of outgrowth was less (Fig. 6D-D”). Importantly, although overall Tubb3 fluorescence was significantly reduced in the anti-N-cadherin-treated explant cultures, reflecting fewer axons observed after blocking N-cadherin-mediated adhesion, there was no change in RFP protein (fluorescence, Fig. 6E) or *RFP* transcript (Fig. 6F) levels between the treatments. These findings reveal that blocking N-cadherin-mediated adhesion did not affect the presence of placodal neurons, but rather impacted axon outgrowth.

**Figure 6.**
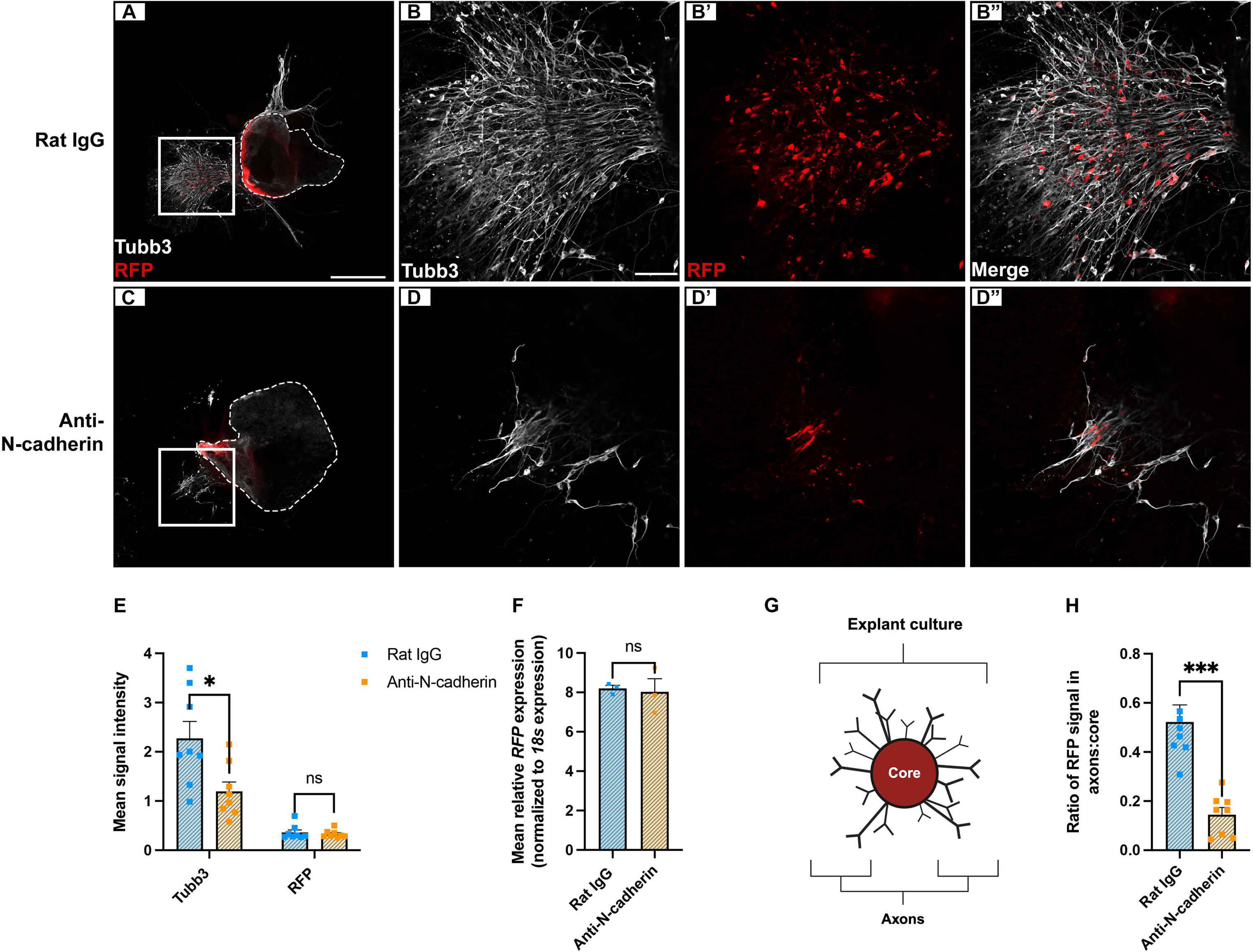
The majority of placodal neurons rely on N-cadherin for axon outgrowth. (A-D”) Representative images of trigeminal explant cultures with RFP-labeled placodal neurons after rat IgG (A-B” (n=8)) or anti-N-cadherin (C-D” (n=8)) treatment and RFP and Tubb3 immunostaining. Boxes in (A) and (C) indicate higher magnification images in (B-B”), and (D-D”), respectively. Dotted lines in (A) and (C) indicate the perimeter of the explanted tissue. Scale bar in (A) is 500 µm and applies to (C); scale bar in (B) is 100 µm and applies to (B’-B”, D-D”). (E) Mean signal intensity for Tubb3 and RFP ± SEM. (F) Quantitative PCR for *RFP* normalized against chick *18s* rRNA of rat IgG- and anti-N-cadherin-treated explants (n=3). (G) Trigeminal ganglion explant culture schematic, with cores (inside the explant) and axons (outside the explant) comprising the explant culture. (H) Ratio of RFP signal found within axons and cores ± SEM. Statistical significance was determined via unpaired t-tests and Mann-Whitney tests (E, F, H) (*) p<0.05, (***) p<0.001.

Given these data and our prior results pointing to a role for N-cadherin in axon outgrowth, we hypothesized that anti-N-cadherin-treated explant cultures would show a greater amount of RFP signal within the core of the explanted tissue, which contains the majority of the neuronal cell bodies, due to the increased presence of placodal neurons unable to extend axons in the absence of N-cadherin-mediated adhesion. To address this, we evaluated the RFP signal found in axonal projections versus RFP signal remaining within the core of the explanted tissue (Fig. 6G). While 33% of the total RFP signal came from axonal projections in rat IgG-treated explants, this dropped to 15% in anti-N-cadherin-treated explants. In contrast, 67% of the total RFP signal was found in the core of the rat IgG-treated explants versus 85% in anti-N-cadherin-treated explants (Fig. 6H). Taken together, these data highlight a cell-autonomous requirement for N-cadherin-mediated adhesion in the proper axon outgrowth of placode-derived neurons.

### Inhibiting N-cadherin-mediated adhesion diminishes neurite complexity in neural crest-derived neurons

Due to the changes we observed in RFP-labeled placodal neurons after blocking N-cadherin-mediated adhesion, we next chose to assess whether neural crest-derived neurons were also impacted. To test this, a neural tube electroporation was performed with PiggyBacGFP with PBase to label the neural crest cells. Next, GFP-positive explant cultures were treated with either rat IgG or anti-N-cadherin as previously described (Fig. 7A), immunostained with a GFP antibody, and neurites were traced (Fig. 7B-C’). Similar to our previous results after N-cadherin MO treatment (Fig. 4), we observed a decrease in neurite complexity after anti-N-cadherin treatment. Neural crest-derived neurons showed a significant reduction in both primary and secondary neurites compared to rat IgG-treated explants (Fig. 7D). In line with our results from Figure 4, we did not observe a change in the average neurite length of primary neurites between treatment groups (Fig. 7E). Very few anti-N-cadherin-treated neural crest-derived neurons had secondary or tertiary neurites, preventing analyses of neurite length. These findings reveal that the adhesive role of N-cadherin impacts the ability of neural crest-derived neurons to generate complex neurites.

**Figure 7:**
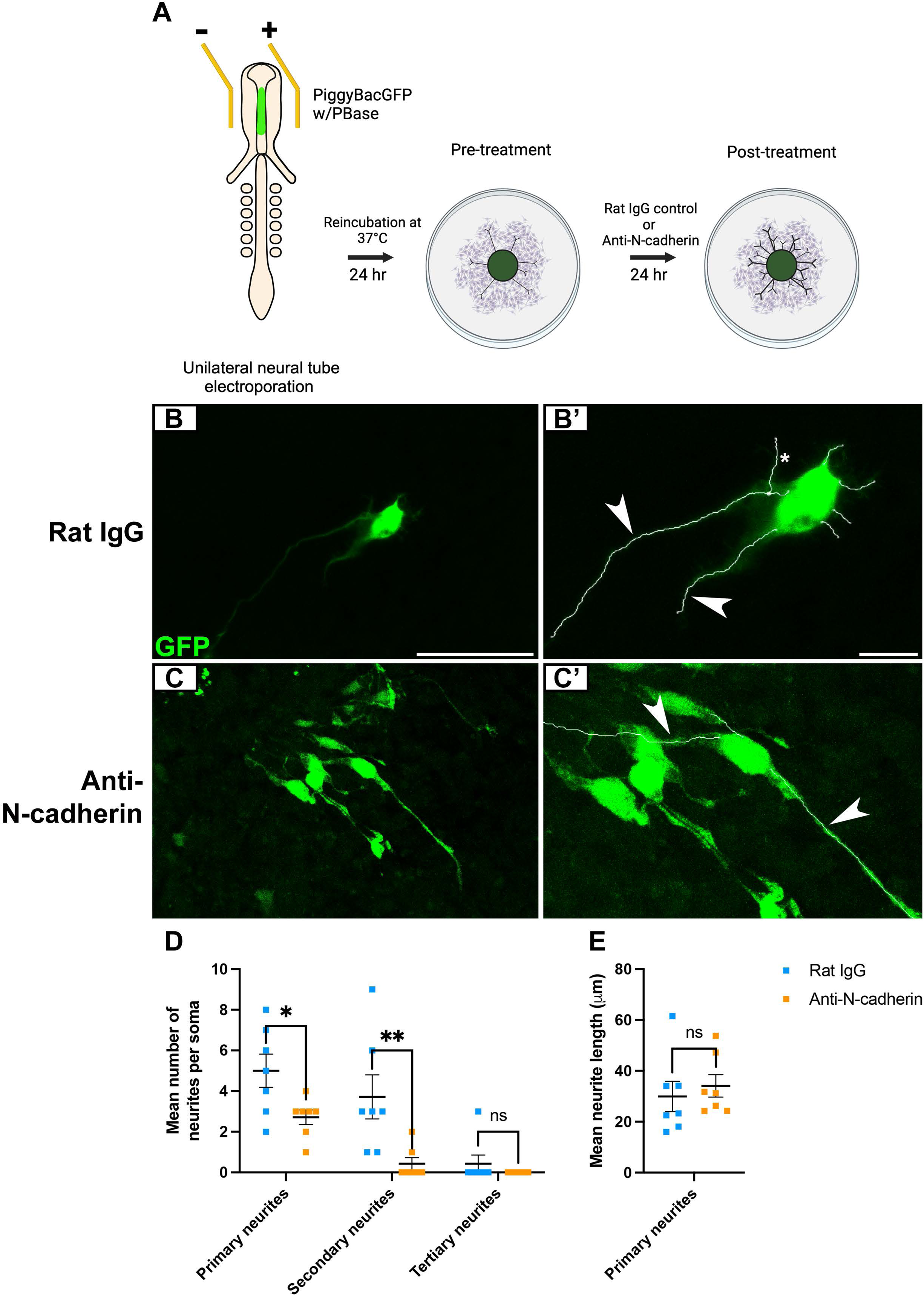
Blocking N-cadherin-mediated adhesion reduces neurite complexity in neural crest-derived neurons. (A) Schematic of N-cadherin function blocking assay after labeling neural crest-derived neurons with PiggyBacGFP. (B-C) Representative images of trigeminal explant cultures with GFP-labeled neural crest-derived neurons after control rat IgG (B, (n=7)) or anti-N-cadherin (C, (n=7)) treatment, GFP immunostaining, and neurite traced images (B’, C’). Arrowheads and asterisks denote examples of primary and secondary neurites, respectively. Scale bar in (B) is 50 µm and applies to (C); scale bar in (B’) is 25 µm and applies to (C’). Mean number (D) and length (E) ± SEM of GFP-positive neural crest-derived neurites after rat IgG or anti-N-cadherin treatment. Statistical significance was determined via unpaired t-tests and Mann-Whitney tests (D-E). (*) p<0.05, (**) p<0.01. Abbreviations: ns: not significant.

## DISCUSSION

Here we demonstrate a sustained role for N-cadherin during trigeminal gangliogenesis, emphasizing the need for cell-cell adhesion in the formation and maturation of the OpV lobe of the trigeminal nerve. Our findings show that reducing N-cadherin from trigeminal placode cells and their neuronal derivatives disrupts trigeminal ganglion morphology and target innervation by specific OpV nerve branches. We found that neural crest-derived neurons rely on N-cadherin in placodal neurons for proper neurite outgrowth and that β-catenin levels are reduced in them after N-cadherin knockdown in placode cells. Moreover, blocking N-cadherin-mediated adhesion led to similar effects, suggesting this underpins the *in vivo* phenotypes observed. These results highlight the continued role of N-cadherin for sensory neurodevelopment and reinforce the importance of reciprocal interactions between placode cells and neural crest cells during cranial ganglia formation.

### Depletion of N-cadherin from placodal neurons leads to persistent morphological defects in the developing trigeminal ganglion

N-cadherin in placode cells plays important roles during early cranial ganglia formation, including facilitating collective cell migration through a repulsive “chase and run” mechanism (Theveneau et al., 2013) and the coalescence of undifferentiated neural crest cells and placodal neurons to generate the trigeminal ganglion (Shiau and Bronner-Fraser, 2009). However, its role in regulating axon extension for proper target innervation has not been explored. Our experiments showed that N-cadherin knockdown initially increased ganglion area, replicating previous results (Shiau and Bronner-Fraser, 2009), but then led to a reduction in trigeminal ganglion size up to E4.5, which eventually recovered. We believe there are two potential reasons for this recovery. First, MOs are typically only stable for up to 4-5 days *in vivo*, reflecting a limitation of our technique (Bill et al., 2009). Therefore, it is possible that at the later timepoints, the MO was targeted for degradation, minimizing phenotypic effects. However, we think this is unlikely, or plays a minimal role, as reduced N-cadherin protein levels were still observed at E6.5/HH28-30. Second, the continued differentiation of neural crest cells into neurons and glia beyond E4-5/HH24-27 (D’Amico-Martel and Noden, 1980) may compensate for this decrease in trigeminal ganglion size, which would not be discernable by Tubb3 immunostaining alone.

In keeping with previous studies, we observed more pronounced phenotypes in OpV compared to MmV, possibly due to the earlier differentiation of OpV neurons compared to MmV cells, which remain proliferative until later stages (Begbie, 2002; Graham and Shimeld, 2013; Lassiter et al., 2014; McCabe et al., 2009; Stark et al., 1997). Although this could be caused by the mosaic nature of the electroporation, we suspect this difference is most likely attributed to MO dilution over time in MmV. OpV placode cells are the first to express neuron-specific markers and become post-mitotic, some even before leaving the ectoderm (McCabe et al., 2009), resulting in more prominent phenotypes. Conversely, MmV placode cells remain as proliferative neuroblasts until after delamination (Begbie, 2002; Graham and Shimeld, 2013; McCabe et al., 2009). N-cadherin depletion from placode cells showed the greatest size difference in OpV at E3.5/HH19-22, coinciding with when the majority of placodal neurons have initiated axon outgrowth (Moody et al., 1989), while no change was observed by E6.5/HH28-30. This suggests that N-cadherin plays a role in promoting axon outgrowth during early stages of neurodevelopment. Notably, we did not observe changes to TrkA- and TrkB-expressing cells when these neurons are prominent (E5.5/HH26-27), indicating altered neuronal differentiation was not a factor in the morphological effects observed. Interestingly, N-cadherin protein levels were still reduced in OpV at E6.5/HH28-30, suggesting continuing effects on N-cadherin even if the MO was targeted for degradation. Additionally, we observed a decrease in cell division in OpV in this stage grouping after N-cadherin knockdown, pointing to other mechanisms that compensate for the absence of phenotype. As only a neuronal marker (Tubb3) was used in these experiments, it is possible that the balance of neural crest cell differentiation into neurons vs. glia was affected, leading to more of the former over the latter.

### Reducing N-cadherin in placodal neurons impacts OpV nerve innervation patterns

Further studies revealed a requirement for N-cadherin in the lateral and medial nasal branches of OpV, with the former exhibiting more drastic changes upon N-cadherin MO treatment. The lateral nasal branch exhibited decreased terminal nerve branches, reduced nerve width, and a smaller innervation area, while the medial nasal nerve showed similar trends for nerve width and innervation area, albeit to a lesser extent. Interestingly, there was no change in the number of terminal nerve branches in the medial nasal branch. This contrasting result - a smaller nerve bundle having similar axonal branching - suggests that N-cadherin functions to adhere individual axons with each other at medial nasal branch targets. Similar phenotypes after N-cadherin perturbations have been observed in other systems as well (Reis et al., 2011; Shikanai et al., 2018; Tan et al., 2010).

Although both OpV branches begin developing at similar stages (Higashiyama and Kuratani, 2014), we chose to examine the E5.5/HH26-27 timepoint for the medial nasal nerve studies due to more consistency among embryos. While differences in embryo age could explain the varying results between the nerve branches, it is equally plausible that the proportion of N-cadherin-expressing placodal neurons differs between them, as the contribution of neural crest- and placode-derived neurons to these nerve branches is unknown. Further, the differences observed could be due to when the respective branches form, as our results suggest that medial nasal branch development occurs after the lateral nasal branch. If true, knockdown of N-cadherin would preferentially impact the lateral nasal nerve since it develops first. Alternatively, the mosaic nature of the electroporation technique may also explain these phenotypes.

### N-cadherin in placodal neurons functions non-cell autonomously to control neurite production in neural crest-derived neurons

Given the observed alterations to trigeminal ganglion morphology, we investigated whether increased apoptosis contributed to these phenotypes. Although our Tubb3 immunohistochemistry results showed no size difference in OpV at the first timepoint examined (E2.5/HH16-18), a significant increase in apoptosis was detected at this timepoint. This suggests that, while the same number of cells may initially contribute to the forming ganglion, some undergo apoptosis, potentially impacting OpV size at later stages and contributing to the long-term phenotypes seen in our studies. However, the absence of any increased apoptosis at later stages, particularly when neural crest cells are differentiating into neurons, implies other mechanisms are also involved.

To address this, we evaluated axon outgrowth of neural crest-derived neurons. Fluorescent labeling of neural crest cells followed by N-cadherin MO treatment in placode cells revealed very few neural crest-derived axons. The complexity of neurites in these neurons was significantly reduced, indicating non-cell autonomous effects. This highlights the continued importance of N-cadherin-mediated interactions throughout trigeminal gangliogenesis, particularly between placode- and neural crest-derived neurons. N-cadherin functions in neurite production for other peripheral neurons (Bixby and Zhang, 1990), and its localization at specific sites along the cell membrane precedes neurite formation (Gärtner et al., 2012). Depletion of N-cadherin from placodal neurons could prevent *trans* N-cadherin interactions between neural crest cells/neural crest-derived neurons and placodal neurons, disrupting neurite production. Additionally, since N-cadherin is also involved in dendrite maintenance (Tan et al., 2010), the stability of neural crest-derived neurites may be compromised when N-cadherin is knocked down in placodal neurons. Interestingly, despite fewer neurites observed in N-cadherin MO-treated explants, neurite length remained the same between treatments, suggesting that processes controlling neurite length are unaffected by knockdown of N-cadherin in placode cells.

Irrespective of the mechanism behind reduced neurite formation, disruptions to the cytoskeleton are likely involved. Our studies revealed a decrease in β-catenin in neural crest-derived neurons upon knockdown of N-cadherin in placode cells. Membrane-bound N-cadherin regulates the availability of both β-catenin and p120-catenin for binding and activating various signaling pathways (Radice, 2013). If *trans* N-cadherin interactions among neural crest- and placode-derived neurons are disrupted, β-catenin could be impacted, leading to the loss of intracellular signaling required to activate GTPases and other cytoskeletal proteins, thus impacting neurite production. Finally, cadherin-receptor tyrosine kinases interactions at cellular membranes reportedly regulate each other (Chiasson-MacKenzie and McClatchey, 2018), including those between N-cadherin and Fibroblast Growth Factor receptors (Hansen et al., 2008; Nguyen and Mège, 2016), which may be critical for the proper growth, maturation, and survival of neural crest-derived neurons. Disrupting N-cadherin levels could interfere with these interactions, affecting the proper development of these neurons.

### N-cadherin-mediated adhesion facilitates axon outgrowth of placode- and neural crest-derived trigeminal neurons

To elucidate whether the *in vivo* phenotypes were due to loss of N-cadherin-dependent adhesion, we used a function-blocking antibody to broadly inhibit N-cadherin in trigeminal ganglion explant cultures. Three main conclusions emerged from the analysis. First, blocking N-cadherin-mediated adhesion did not preclude axon outgrowth. Second, intersection analyses revealed reduced axon outgrowth and branching in anti-N-cadherin-treated explants compared to rat IgG-treated controls. However, anti-N-cadherin-treated explants still extended axons to the periphery, though not significantly more than pre-treatment explants, suggesting some axons may be retracting, due to either pruning or lack of necessary factors in the media to support axon maintenance. Lastly, segment lengths (used as a proxy for axons) were significantly reduced in our rat IgG-treated explants compared to both pre-treatment and anti-N-cadherin-treated explants. This reduction was due to a higher number of very short segments in the rat IgG group, as longer axon lengths were observed in all groups and argues against a substantial role for axon pruning. Since some trigeminal neurons could extend axons similarly to controls despite N-cadherin inhibition, we conclude that N-cadherin is not essential for axon outgrowth in all trigeminal neurons. These data align with the *in vivo* observation for OpV branches in that N-cadherin knockdown does not completely block target innervation but reduces the number of axons reaching their targets. While this phenotype could be attributed to the mosaic nature of the MO electroporation, our *in vitro* results now offer another potential explanation, in that some neurons are not reliant on N-cadherin for the outgrowth of their axons.

We next identified the cellular origin of the neurons requiring N-cadherin-mediated adhesion for their outgrowth, hypothesizing that they would be placode-derived since placode cell neuronal differentiation precedes neural crest cell differentiation *in vivo* (D’Amico-Martel and Noden, 1983). Our experiments showed reduced axon outgrowth in RFP-labeled placodal neurons after anti-N-cadherin-treatment. Although some neurons might have initially extended axons and then died, we believe this is unlikely because a subset of neurons was always capable of growing axons irrespective of when the anti-N-cadherin antibody was added to the explants. Although RFP transcript and protein levels were similar between treatments, very interestingly, RFP signal in axonal projections was reduced after anti-N-cadherin treatment, with most RFP signal originating from inside the cores. This indicates that N-cadherin promotes axon outgrowth, with its absence limiting the number of RFP-positive placodal neurons capable of extending axons.

Although we did not observe interactions between fluorescently labeled placodal neurons and neural crest-derived neurons *in vitro* upon blocking N-cadherin-mediated adhesion, we did see changes in neurite production of neural crest-derived neurons. While both primary and secondary neurite number was significantly reduced, neurite length remained unchanged across treatments. These results mirrored phenotypes in neural crest-derived neurons when N-cadherin was knocked down from placode cells, indicating that the loss of N-cadherin from placodal neurons impacts the ability of neural crest-derived neurons to produce neurites. This likely contributes to the reduced size of the trigeminal ganglion observed in N-cadherin knockdown embryos, as these neurons may be unable to extend axons to their targets properly.

Surprisingly, a small subset of RFP-positive placodal neurons exhibited axon outgrowth after anti-N-cadherin treatment, potentially representing pioneer neurons of the trigeminal ganglion. Though pioneer neurons are well-documented in invertebrate models (Clarke et al., 2020; Hidalgo and Brand, 1997; Klose and Bentley, 1989; Lin et al., 1995), evidence in vertebrates (Eisen et al., 1986; Ghosh et al., 1990; Gong and Shipley, 1995), including in mouse trigeminal neurodevelopment, suggests that the first axons to extend from OpV provide a path for secondary axons to reach their targets (Stainier and Gilbert, 1990). A similar phenomenon has been observed in the chick trigeminal ganglion, where a few placode-derived neurons were observed to extend axons to the periphery, providing a path for later-born neurons to follow (Moody et al., 1989). Collectively, these studies and our new data suggest that a subset of placodal neurons may serve as pioneers upon which other placode-derived neurons and later neural crest-derived neurons (“followers”) extend axons (Moody et al., 1989). This N-cadherin-independent (pioneer) and -dependent (follower) outgrowth is reminiscent of mechanisms regulating migration in the facial branchiomotor neuron system (Wanner and Prince, 2013).

## CONCLUSION

In this study, we have shown a sustained negative impact on the trigeminal ganglion following N-cadherin depletion from placodal neurons. Throughout development, including after neural crest cells differentiate into neurons, we observed changes in trigeminal ganglion size and reduced axon outgrowth to target tissues. These morphological differences could not be fully explained by increased apoptosis. In support of this, neurite production and β-catenin levels in neural crest-derived neurons were altered after depleting N-cadherin from placode cells. Treatment of explants with an N-cadherin adhesion-blocking antibody caused similar effects on neurites, pointing to a non-cell autonomous requirement for N-cadherin in placodal neurons to facilitate axon outgrowth from placode- and neural crest-derived trigeminal neurons. Inhibiting N-cadherin-mediated adhesion *in vitro* also reduced axon outgrowth from most placodal neurons, demonstrating a cell autonomous function for N-cadherin in this process. Altogether, our results add to the growing amount of evidence supporting the importance of reciprocal interactions between neural crest cells and placode cells during the formation of the cranial ganglia, and highlight a long-term, critical role for N-cadherin during trigeminal gangliogenesis that will likely be translatable to other cranial ganglia and vertebrate species.

## MATERIALS AND METHODS

### Chick embryos

Fertilized chicken eggs (*Gallus gallus*) were obtained from the Department of Animal and Avian Sciences at the University of Maryland (MD) and Centurion Poultry Incorporated (GA) and incubated at 37°C in humidified incubators on their sides. Embryos were incubated to reach desired Hamburger Hamilton (HH) stages based on times provided therein (Hamburger and Hamilton, 1992).

### Morpholinos

To reduce N-cadherin protein expression, a validated lissamine-tagged translation-blocking morpholino antisense oligonucleotide (MO) against the *N-cadherin* transcript was used (GeneTools, LLC) (Shiau and Bronner-Fraser, 2009), with a sequence of 5’-GCGGCGTTCCCGCTATCCGGCACAT-3’. For some control experiments, a 5 base pair mismatch N-cadherin control MO (N-cadherin MM) was used (Shiau and Bronner-Fraser, 2009), with a sequence of 5’-GCGcCcTTCCCcCTATgCcGCACAT-3’ (altered base pairs in lowercase). Both MOs were diluted with ultrapure water and 10% sucrose (in 1X phosphate-buffered saline (PBS)) and used at a final concentration of 500 µM.

### *In ovo* unilateral placode electroporation

To target trigeminal placode cells prior to their delamination, a unilateral electroporation was performed at E1.75/HH10-11. The N-cadherin MO, N-cadherin MM, or a combination of PiggyBacRFP, a genome incorporating vector (Chen and LoTurco, 2012) (0.819 µg/µl), with PBase, its transposase (0.912 µg/µl), was expelled on top of the ectoderm on one side of the embryo using a fine glass needle. Next, a 30-gauge needle was used to puncture a small hole through the vitelline membrane so that electrodes could lay vertically above and below the ectoderm over which the MO/plasmid had been placed. Three, 9V pulses were delivered over 50 ms at 200 ms intervals. After electroporation, eggs were resealed with tape and parafilm and grown at 37°C to desired stages of interest. Due to the inherent variations that arise during normal development, even when embryos are staged identically, the contralateral side of the electroporated embryo was a more appropriate control and was used for all *in vivo* studies. Embryos were screened for successful MO electroporation based on the red fluorescence of the MO prior to collection and use in subsequent assays.

### Sequential electroporation method

To target both cell populations that contribute to the trigeminal ganglion, a modified sequential electroporation (Halmi et al., 2022) was utilized. Briefly, a unilateral neural tube co-electroporation was first performed on E1.5/HH9 using a combination of PiggyBacGFP (0.819 µg/µl) and PBase (Wang et al., 2008) (0.912 µg/µl). Two, 15V pulses were delivered over 30 ms at 200 ms intervals. The electroporated embryos were re-incubated until they reached E1.75/HH10-11, and then the placode cells on the same side of the embryo were electroporated with the N-cadherin MO, N-cadherin MM, or PiggyBacRFP in the same manner as described above. After this second electroporation, eggs were resealed with tape and parafilm and grown at 37°C to desired stages of interest.

### Immunostaining

Electroporated embryos were collected at desired stages of interest and fixed in 4% paraformaldehyde (PFA) overnight at 4°C with agitation. Following fixation, embryos were washed 3 x 10 minutes in 1X PBS and stored in 1X PBS at 4°C until further processing. Primary antibodies used included Tubb3 (Abcam ab78078, 1:500 for section, 1:250 for whole-mount), GFP (Abcam ab6662, 1:250), RFP (Invitrogen MA5-15257, 1:250), TrkA (gift from Dr. Francis Lefcort, Montana State University, 1:5000), TrkB (gift from Dr. Francis Lefcort, Montana State University, 1:2500), cleaved caspase-3 (Cell Signaling Technology #9664, 1:100), β-catenin (Santa Cruz Biotechnology sc-393501 1:100). Secondary antibodies used at 1:250 included goat anti-rabbit IgG 488 (Invitrogen A211034), goat anti-mouse IgG1 594 (Invitrogen A21125), goat anti-mouse IgG2b 647 (Invitrogen A21242), goat anti-mouse 405 (Invitrogen A31553), and goat anti-mouse IgG2a 555 or 647 (SouthernBiotech 1080-31 or 1080-32).

#### Tissue sections

Embryos were processed through an increasing sucrose gradient (5% and 15% sucrose (Fisher) in 1X PBS) at 4°C either overnight or until they sank to the bottom of the glass vials in which they were stored. Next, they were processed through 7.5% gelatin (Sigma) in 15% sucrose and 20% gelatin in 1X PBS, both at 37°C, either overnight or until the embryos sank. Embryos were embedded in 20% gelatin, flash-frozen using liquid nitrogen, and stored at -80°C until ready for use. Embedded embryos were then cryo-sectioned at 12 µm on charged slides (VWR, 48311-703). Slide-mounted tissue sections were stored at -20°C until ready for use. For immunohistochemistry, slides were de-gelatinized by incubation in 1X PBS at 42°C for 15-20 minutes. Following 3 x 5-minute washes in 1X PBS, the slides were blocked with 10% heat-treated sheep serum (Sigma, HTSS) in 1X PBS + 0.1% Triton X-100 (Sigma, PBST) at room temperature for 1-2 hours in a humidified chamber, then rinsed in PBST. Primary antibodies were diluted in PBST + 5% HTSS, added to the slides, covered with a glass coverslip (Fisher) to prevent sections from drying out, and incubated overnight at 4°C. The following day, the coverslips were removed, and the slides were washed 4 x 30 minutes in PBST. Secondary antibodies were then diluted in PBST + 5% HTSS, added to the slides, covered with a glass coverslip, and incubated overnight at 4°C. The next day, slides were again washed 4 x 30 minutes in PBST and mounted with DAPI-fluoromount media (SouthernBiotech).

#### Standard whole-mount immunohistochemistry

Embryos were blocked with 10% HTSS in PBST for 2 hours at room temperature. Following a rinse with PBST, primary antibodies were diluted in PBST + 5% HTSS and incubated overnight at 4°C with agitation. The following day, the embryos were washed 4 x 30 minutes in PBST. Secondary antibodies were diluted in PBST + 5% HTSS and incubated overnight at 4°C with agitation. The next day, embryos were again washed 4 x 30 minutes in PBST and stored in 1X PBS at 4°C until they were either imaged or cleared (see below).

#### Modified whole-mount immunohistochemistry for iDISCO clearing

An adapted protocol was used based on methods from (Renier et al., 2014). Fixed embryos were washed twice for 1 hour in 1X PBS and were then taken through an increasing methanol (MeOH, Pharmco) in 1X PBS gradient (50% (1 x 1 hour), 80% (1 x 1 hour), 100% (2 x 1 hour)), before they were bleached overnight in 5% H2O2 (Sigma) in 20% DMSO (Fisher)/MeOH at 4°C. The next day, samples were taken through the following series of washes: 2 x 1 hour in 100% MeOH, 2 x 1 hour in 20% DMSO/MeOH, 1 x 1 hour in 80% MeOH, 1 x 1 hour in 50% MeOH, 2 x 1 hour in 1X PBS, and 2 x 1 hour in 1X PBS + 0.2% Triton X-100. Samples were incubated overnight in 20% DMSO/0.3M glycine (Fisher) in 1X PBS + 0.2% Triton X-100 at 4°C. The next day, samples were blocked in 10% HTSS/10% DMSO/1X PBS + 0.2% Triton X-100 overnight at 4°C. Embryos were washed 2 x 1 hour in 0.2% Tween-20 (Sigma) in 1X PBS with 10 µg/ml heparin (Fisher, PTwH) at room temperature before primary antibodies were added and incubated for 72 hours at 4°C. Primary antibodies were diluted in 5% DMSO/5% HTSS/PTwH. Samples were washed in PTwH for 24 hours before secondary antibodies were added (diluted in the same manner as the primary antibodies), and samples were incubated with secondary antibodies for 48 hours at 4°C. Samples were then washed for 48 hours in PTwH before iDISCO clearing (see below).

#### Explant immunocytochemistry

Explant slides were washed once with 1X PBS, followed by fixation in 4% PFA for 1 hour at room temperature with agitation. Following fixation, slides were rinsed with 1X PBS and stored in 1X PBS at 4°C until ready for immunocytochemistry. A hydrophobic perimeter was drawn around the slides to prevent dehydration of tissue using an ImmEdge Pen (Vector Laboratories), and explants were taken through blocking, primary antibody incubation, and secondary antibody incubation in the absence of coverslips as described in the tissue section methods. Slides were mounted with DAPI-fluoromount media (SouthernBiotech) and covered with a glass coverslip.

### Tissue clearing

As embryos progress through development and tissue structures surrounding the trigeminal ganglion become more complex, visualization of the trigeminal ganglion after whole-mount immunohistochemistry becomes increasingly difficult. Therefore, clearing methods to render the tissues transparent were performed. At early stages (up to E3.5/HH22), the FRUIT (Hou et al., 2015) tissue clearing protocol was used, whereas later stages required iDISCO (Renier et al., 2014) tissue clearing. With both protocols, MO fluorescence was quenched during the clearing process.

#### FRUIT

Following the standard whole-mount immunohistochemistry protocol, embryos were progressed through increasing concentrations of fructose (35%, 40%, 60% and 80% w/v) in 8M urea (Sigma), 0.5% (v/v) alpha-thioglycerol (Fisher) dissolved in ultrapure water (“FRUIT buffer”) for a minimum of 8 hours, up to 24 hours at room temperature, depending on the stage of the embryo. A sample was only moved up to the next FRUIT buffer once it had visibly changed transparency. The FRUIT cocktail, containing increasing concentrations of fructose, was made by first dissolving fructose in ultrapure water at 37°C. Once dissolved, the urea was added, and the solution was rocked at room temperature until fully dissolved. Lastly, alpha-thioglycerol was added, and buffers were stored at 4°C until use. Embryos were imaged in 60% or 80% FRUIT buffer (depending on the stage of the embryo) in glass-bottomed petri dishes (MatTek P35G-1.5-20-C). Typically, embryos up to E3/HH19 were imaged in 60% FRUIT while older embryos were imaged in 80% FRUIT.

#### iDISCO

Following the modified whole-mount immunohistochemistry protocol, embryos were washed in 1X PBS + 0.2% Triton X-100 for a minimum of 48 hours. Samples were incubated in 50% (v/v) tetrahydrofuran (Sigma, THF) in water overnight in glass vials, making sure that over 90% of the vial was filled with the solution. The following day, embryos were then incubated in 80% THF in water for 1 hour, followed by 2 x 1 hour incubations in 100% THF. Next, samples were incubated in 100% dichloromethane (Sigma) for 1-2 hours (or until they sank to the bottom of the vial). Lastly, embryos were incubated in dibenzyl ether (Sigma, DBE) overnight and imaged the following day in DBE. All steps, including embryo storage in DBE, were performed at room temperature. iDISCO-cleared embryos were imaged in a glass petri dish containing a custom chamber (Renier et al., 2014), which was printed in Terrapin Works, part of the School of Engineering at the University of Maryland.

### TUNEL assay

To assess apoptosis (TUNEL), the *In Situ* Cell Death Detection Kit, Fluorescein (Roche) was used according to manufacturer’s instructions. Briefly, following secondary antibody washes (4 x 30 minutes in PBST), slides were washed once for 30 minutes in 1X PBS. Slides were re-fixed with 4% PFA for 20 minutes at room temperature, followed by one 30-minute wash in 1X PBS. The TUNEL reaction mixture was added to the slides (75 µL per slide), and a glass coverslip was placed over them. Slides were incubated in the dark for 1 hour at 37°C. Next, the coverslips were removed, and the slides were washed 3 x 5 minutes in 1X PBS. Lastly, slides were mounted with DAPI-fluoromount media (SouthernBiotech).

### Trigeminal ganglion explant assay

Wildtype E2/HH13-14 embryos were removed from eggs and stored on ice in 1X PBS until dissections. Trigeminal ganglia were dissected on a sylgard-coated dish by first bisecting the embryo and then carefully taking the area of the forming trigeminal ganglion, making sure to exclude the eye and neural tube. Explants were cultured on 2-well glass chamber slides (ThermoFisher) coated with Poly-L-Lysine (0.01% PLL, Sigma) and fibronectin (0.01%, Sigma). To make the coated slides, PLL was added to each chamber on the glass slides and incubated inside a biosafety cabinet for 30 minutes at room temperature. The PLL was removed, and slides were washed 3 times with water. Next, fibronectin (diluted in DMEM media (ThermoFisher)) was added to each chamber and incubated at 37°C for a minimum of 2 hours. The fibronectin solution was removed and slides were stored at 4°C until needed (for up to one week). Explants were grown in DMEM media supplemented with N2 (1%, ThermoFisher) and Penicillin/Streptomycin (1%, Fisher). Explants for N-cadherin function-blocking antibody experiments (see below) were grown in the supplemented DMEM media mentioned above. Sequentially-labeled cells for N-cadherin MO- or MM-containing explant experiments were grown in Neurobasal media (ThermoFisher) supplemented with B27 (1%, ThermoFisher), N2 (0.5%, ThermoFisher), Glutamax (1%, ThermoFisher), Penicillin/Streptomycin (1%, ThermoFisher), and with nerve growth factor (ThermoFisher), brain-derived neurotrophic factor (Amgen), and neurotrophin-3 (Amgen), all at 50 ng/ml.

### N-cadherin function-blocking antibody experiments

After 24 hours of incubation, fresh media containing either an antibody to block N-cadherin function, MNCD2 (anti-N-cadherin, Developmental Studies Hybridoma Bank, 0.1 µg/ml), or a normal rat IgG control (Santa Cruz Biotechnology, 1.6 µg/ml), replaced the initial explant plating media, and explants were incubated overnight. The following morning, the slides were briefly rinsed with 1X PBS and then fixed in 4% PFA for one hour at room temperature. The explants were washed 3 times in 1X PBS and then processed for immunocytochemistry.

### Immunoblotting

E6.5/HH28-30 N-cadherin MO-electroporated and contralateral OpV lobes from six embryos per replicate were dissected, pooled, and lysed in lysis buffer (50 mM Tris pH 7.5, 100 mM NaCl, 0.5% IGEPAL CA-630) supplemented with cOmplete Protease Inhibitor Cocktail (Roche) and 1 mM PMSF for 30 minutes on ice with periodic gentle vortexing. Lysates were then centrifuged at maximum *g* for 20 minutes at 4°C to separate fractions and the soluble protein fraction was then quantified by Bradford assay (ThermoFisher). 12 µg of each sample was boiled at 99°C for 5 minutes in 4x reducing Laemmli sample buffer. Samples were separated by 12% SDS-PAGE then transferred to PVDF membrane (ThermoFisher). Membrane was blocked in 5% milk (in 1X PBS + 0.1% Tween-20) for 30 minutes at room temperature followed by overnight incubation at 4°C with the following primary antibodies diluted in blocking solution: Phospho-histone H3 (Sigma 06570, 1:500), N-cadherin (DSHB MNCD2, 1:100), and GAPDH (GeneTex GTX627408 1:1000). Membranes were washed 3 x 10 minutes in 1X PBS + 0.1% Tween-20 before incubation with secondary antibodies (species- and isotype-specific horseradish peroxidase, Jackson ImmunoResearch) diluted in blocking solution for 1 hour at room temperature. Membranes were washed 3 x 10 minutes in 1X PBS + 0.1% Tween-20 and antibody detection was performed using Supersignal West Pico or Femto chemiluminescent substrates (ThermoFisher) and visualized using a ChemiDox XRS system (Bio-Rad).

### Quantitative PCR

Trigeminal ganglion explants in which placode cells were electroporated with PiggyBacRFP plus PBase underwent rat IgG or anti-N-cadherin treatment (all as described above). Explant cultures were pooled and lysed in RNA lysis buffer and total RNA was extracted using the RNase Mini Kit (Qiagen). cDNAs were synthesized using the SuperScript IV One-Step RT-PCR System (ThermoFisher) according to manufacturer’s instructions. Quantitative PCR was performed with iTaq Universal SYBR Green Supermix (BioRad) according to manufacturer’s instructions. Equal amounts of cDNA for rat IgG- and anti-N-cadherin-treated samples were used for quadruplicate reactions and mixed with SYBR Green Supermix, and 750 nM of each primer (RFP or chick 18s rRNA). To validate results, quantitative PCR was also performed on cDNA samples prepared in the absence of reverse transcriptase and any template cDNA (no template control). *RFP* levels were normalized against chick *18s* rRNA levels, as described previously (Schiffmacher et al., 2016). Average normalized values of RFP were plotted as bar graphs. Primer sequences are as follows: RFP Forward: (5’-CTCCGAGGACGTCATCAA-3’), RFP Reverse: (5’-TTGGTCACCTTCAGCTTGG-3’), chick *18s* rRNA Forward: (5’-TGTGCCGCTAGAGGTGAAATT-3’), and chick 18s rRNA Reverse: (5’-TGGCAAATGCTTTCGCTTT-3’).

### Confocal imaging

All images were acquired with the LSM Zeiss 800 confocal microscope (Carl Zeiss Microscopy, Thornwood, NY, USA). Laser power, gain, and offset were kept consistent for the different channels during all experiments where possible. Fiji (NIH) (Schindelin et al., 2012) and Adobe Photoshop CC 2023 were used for image processing. For whole-mount imaging with Z-stacks, all embryos were imaged at a 5 μm interval. For sections, images of at least five serial transverse sections were acquired and used for analysis. For all experiments, a minimum of five replicates was obtained.

### Quantifications and statistical analyses

All quantifications were done on Fiji and all graphs were generated using Prism GraphPad Version 10.

#### Area measurement

Maximum intensity Z-projections were generated through the trigeminal ganglion. The polygon tool was used to outline the perimeter of Tubb3 signal, and area measurements were taken for the whole trigeminal ganglion, the OpV lobe, and the MmV lobe. Individual area measurements for each developmental stage grouping were averaged together and plotted to generate graphs.

#### Cell counting

Cell counts were either performed manually or using the Squassh (Rizk et al., 2014) plugin from the MOSAICsuite (Shivanandan et al., 2013), as described (Halmi et al., 2022). The Tubb3 channel of each image was used to crop the trigeminal ganglion, which was then used to crop the images for the channel of interest. For Squassh, segmentation parameters were kept uniform for all images and run as a batch function. The quality of image segmentation was confirmed visually, and adjustments were made as necessary. The number of puncta/cells was calculated per image and averaged to obtain a mean number per replicate, which was then divided by the average area of the trigeminal ganglion to adjust for size differences. Individual averages for each developmental stage grouping were plotted to generate graphs.

#### Nerve branching

Z-stack images of the OpV branch of the trigeminal nerve were analyzed using custom ImageJ Macro language (IJM) scripts (available upon request). Maximum intensity Z-projections were generated and input into the ImageJ macro. Signal profiles were obtained via successive vertical line selections using *getProfile()* at 30 μm intervals. The number of terminal nerve branches present at increasing distances from the starting point were exported and calculated in R (R core team, Vienna, Austria, 2021, scripts available upon request). Data were aligned to ensure consistent starting points for each replicate, and the mean number of nerve branches was plotted to generate graphs.

#### Nerve width

Nerve width measurements for both the lateral and medial nasal nerves were generated using maximum intensity Z-projections and the straight line tool. The width of individual OpV nerve divisions (prior to extensive branching within the target tissue) was measured based on the Tubb3 signal for each embryo. Widths for N-cadherin MO-treated embryos and their contralateral control sides were averaged and plotted to generate graphs.

#### Innervation area

Maximum intensity Z-projections were used to generate innervation area measurements for both the lateral and medial nasal nerves. The polygon tool was used to draw a border around the Tubb3 signal in the target tissues where terminal nerve branching occurred. Area measurements for N-cadherin MO-treated embryos and their contralateral control sides were averaged and plotted to generate graphs.

#### Intersection analysis following treatment with anti-N-cadherin

Background was removed and the ridge-detection plug-in (Steger, 1998) was used to highlight axonal outgrowth and to quantify axon length from explant images. Using the detected segment binary images and the concentric circle plugin, 20 concentric circles were overlayed on the explant, centered at the middle. The number of axonal intersections at each successive ring of increasing diameter was counted and plotted to generate graphs.

#### RFP-labeled placode cell analysis

Background was eliminated from explant images of the Tubb3 channel and the polygon tool was used to isolate the core of the explant from its associated axonal projections. The fluorescent intensity of RFP signal was collected for both the axonal projections and the core of the explants. The mean axonal signal was divided by the mean core signal to generate a ratio. Ratios for individual treatments were averaged and compared between conditions.

#### Simple neurite tracer analysis

Z-stacks of GFP-positive neural crest-derived neurons were analyzed using the simple neurite tracer plugin (Arshadi et al., 2021; Ferreira et al., 2014). Primary neurites were traced from the cell body through the entire Z-stack. Neurites extending from primary neurites were added as branch points and termed secondary neurites. Any neurite branching off secondary neurites was marked as a tertiary neurite. The mean number and mean length of primary, secondary, and tertiary neurites were plotted to generate graphs. **Immunoblot band intensity:** Immunoblot images were processed with the Analyze Gel function. Equal boxes were drawn around the lanes of interest using the rectangle tool, and band intensities were plotted as peaks of signal. The line tool was used to isolate peaks and remove background, measuring the area under the peak to quantify band intensity for each protein. Phospho-histone H3 and N-cadherin were normalized against GAPDH and plotted to generate graphs.

#### β-catenin analysis

Z-stack images of sequentially-electroporated explants were processed using custom ImageJ Macro language (IJM) scripts (available upon request). GFP-positive neurons were identified over each slice using thresholding. The area of each identified cell was calculated using the analyze particles tool, and the mean β-catenin signal was quantified from the overlapping GFP signal in the Z-stacks. Slices with insufficient β-catenin staining were excluded. Mean β-catenin signal was plotted to generate graphs.

## ACKNOWLEDGMENTS

The authors would like to thank the National Institute of Dental and Craniofacial Research (R01DE024217) and the University of Maryland Graduate School for financially supporting this research. Additionally, we would like to thank Ms. Zoe Davidson and Ms. Reagan Armstrong for their technical assistance. All schematics were created with Biorender.com.

## DATA AND RESOURCE AVAILABILITY

All relevant data can be found within this article and the supplementary information.

## SUPPLEMENTAL FIGURE LEGENDS

**Supplemental Figure 1. N-cadherin mismatch control MO (MM)-treated embryos show normal trigeminal ganglion morphology.** (A-A’) Representative images of the trigeminal ganglion after Tubb3 immunostaining (E4.5/HH23-25, (n=4)). Scale bar in (A) is 250 µm and applies to (A’). (B) Mean whole trigeminal ganglia area ± SEM for contralateral control and N-cadherin MM sides of embryos. Statistical significance was determined via an unpaired t-test. Abbreviations: TG: trigeminal ganglion; OpV: ophthalmic; MmV: maxillomandibular; MM: mismatch control; MO: morpholino; ns: not significant.

**Supplemental Figure 2: N-cadherin knockdown in placode cells does not increase cell proliferation at later developmental stages.** (A) Immunoblots of E6.5/HH28-30 electroporated (MO) and contralateral (CNTL) trigeminal OpV lobes for N-cadherin, phospho-histone H3 (PHH3), and GAPDH (n=2). (B) Protein levels of N-cadherin and phospho-histone H3 normalized against GAPDH loading control, presented as a fraction of total signal from contralateral control OpV lobes.

**Supplemental Figure 3: N-cadherin depletion from placode cells does not alter subpopulations of sensory neurons within the trigeminal ganglion.** (A-D’) Sagittal serial sections through the OpV lobe of the trigeminal ganglion (contralateral control (A-B’) and N-cadherin MO-electroporated side (C-D’)) from representative E5.5/HH26-27 (n=4) embryos with TrkA and TrkB immunostaining. Images of TrkA and TrkB serial sections were merged to generate representative images. Scale bar in (A) is 100 µm and applies to (C); scale bar in (B) is 50 µm and applies to (B, D-D’). (E) Average number of TrkA- or TrkB-positive OpV neurons ± SEM after N-cadherin MO electroporation. (F) Average number of TrkA- or TrkB-positive OpV neurons normalized against OpV lobe area ± SEM. Statistical significance was determined via paired t-tests (E, F). (*) p<0.05.

**Supplemental Figure 4. Sequential electroporation of neural crest cells and placode cells exclusively labels distinct cell populations**. Representative images of an HH25/E4.5 embryo sequentially electroporated with PiggyBacGFP plus PBase (neural crest cells) and N-cadherin MM (placode cells), followed by Tubb3 and GFP immunostaining. Scale bar in (A) is 100 µm and scale bar in (B) is 50 µm and applies to (B’-B’’).

**Supplemental Figure 5. N-cadherin MO-treated explants qualitatively exhibit less neural crest-derived axon outgrowth.** (A-B’) Representative images of sequentially-electroporated trigeminal explants with neural crest cells co-electroporated with PiggyBacGFP plus PBase, and placode cells electroporated with N-cadherin MM (A-A’) or N-cadherin MO (B-B’), followed by GFP and Tubb3 immunostaining. Arrowheads in (A’) and (B’) point to neural crest-derived axons. Scale bar in (A) is 100 µm and applies to all images.

