## Supplementary figures and images for "N-cadherin facilitates trigeminal sensory neuron outgrowth and target tissue innervation"

### Supplemental Figures

Figure 1:

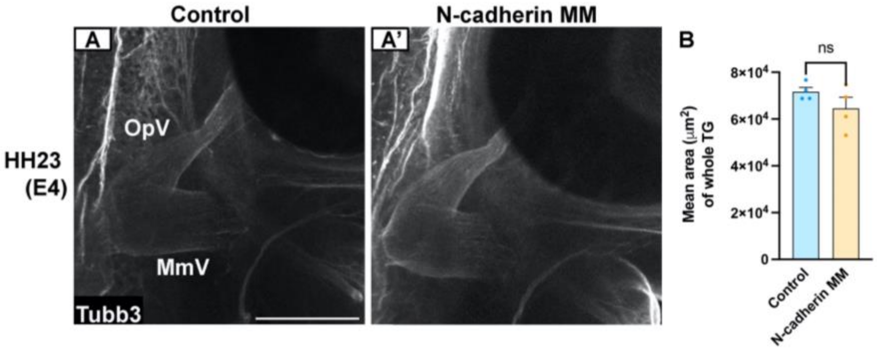


Figure 2:

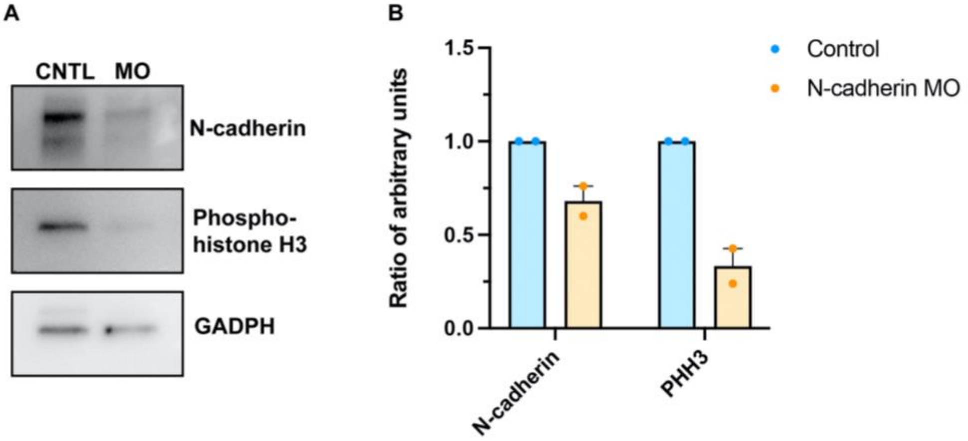

Figure 3:

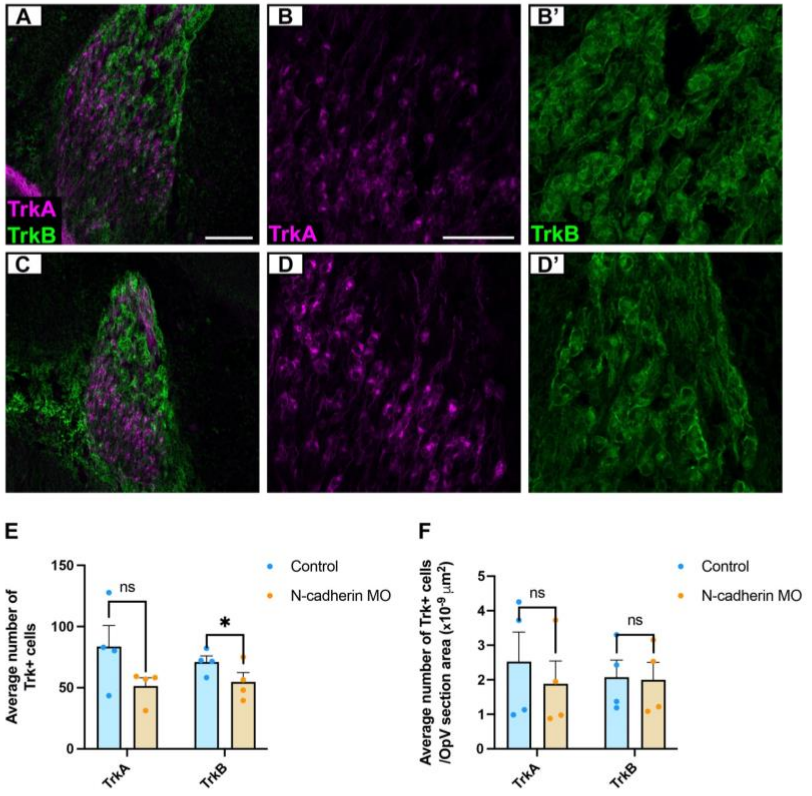


Figure 4:

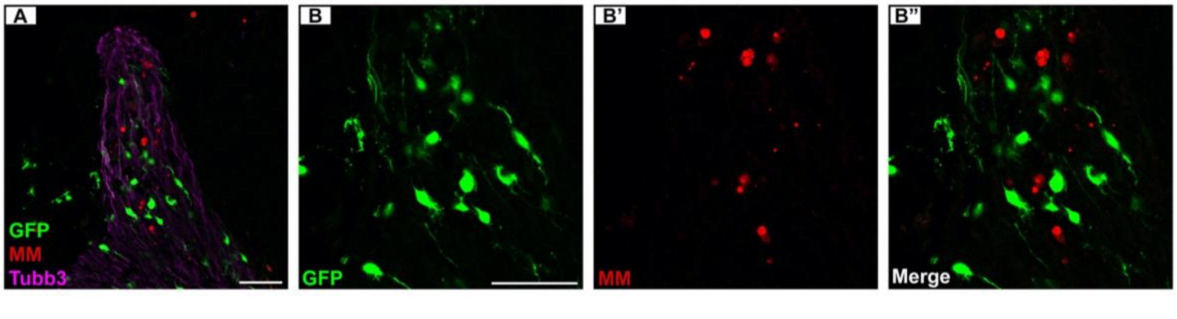

Figure 5:

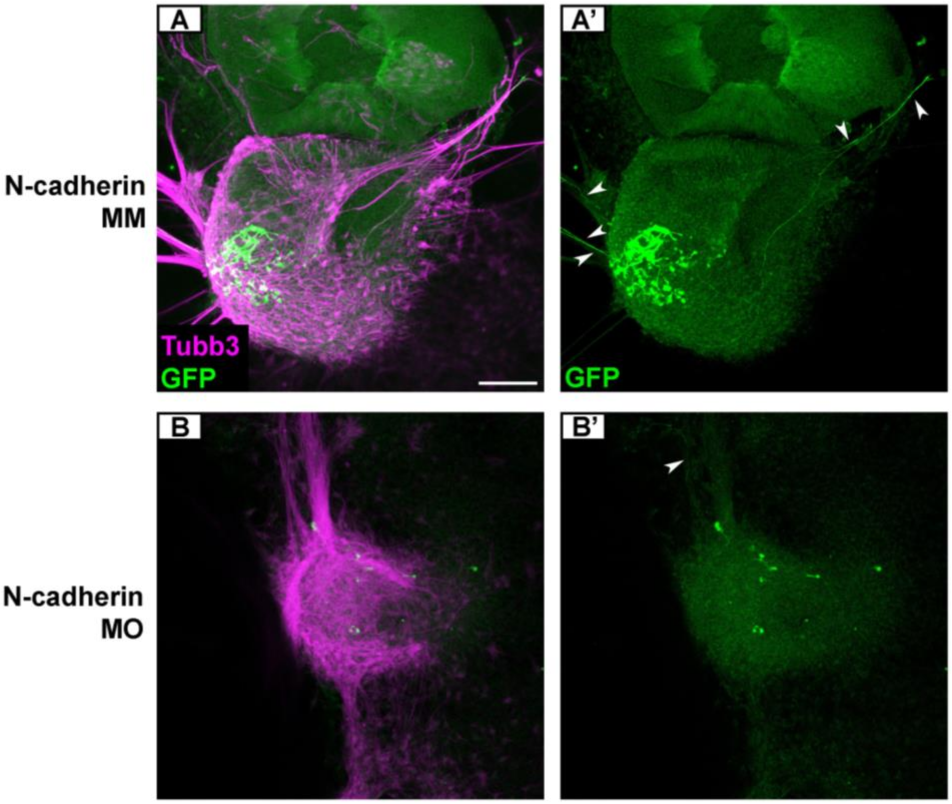
